# Exploring the likelihood of the membrane pollution hypothesis with a physiologically plausible biophysical model

**DOI:** 10.1101/2025.10.06.680798

**Authors:** José Luis Hernández-Cáceres, Liudmila Lukianenko, Elena Venskaya, Yury D. Nechipurenko, Yury L. Orlov, Jorge Iglesias-Fuster, Lidice Galán-García

## Abstract

Epilepsy affects more than 52 million people worldwide and has been known since ancient times. Despite this long history, available therapeutic methods—both pharmacological and non-pharmacological—fail to control seizures in over 10% of patients. At the same time, there exists a diversity of theories regarding the fundamental mechanisms underlying epilepsy. Understanding the biophysical basis of the simplest manifestation of epileptic activity—the paroxysmal depolarization shift (PDS)—could therefore be highly valuable.

Prevailing ideas consider PDS as exaggerated synaptic excitatory potentials, but experimental evidence shows that PDS can be generated in isolated neurons devoid of synaptic input. Ulrich Altrup proposed an alternative view, suggesting that PDS represent aberrantly large pacemaker potentials rather than giant excitatory postsynaptic potentials. Theoretical work by Hernandez-Caceres and Brenes supported this concept, demonstrating a bifurcation-like transformation from physiological pacemaker potentials to PDS and finally to long-duration sustained depolarizations.

According to Altrup’s membrane pollution hypothesis (MPH), epileptic activity emerges as a consequence of incorporating amphiphilic pollutants into neuronal membranes, which leads to an increase in membrane micro-viscosity. Using a biophysically sound model for pacemaker activity, we explored the possibility of eliciting PDS through increased membrane micro-viscosity. Provided results suggest that this is theoretically plausible.

Further experimental and theoretical research is needed to refine details of the MPH and to develop new strategies for combating epilepsy.

## Introduction

About 52 million persons worldwide suffer from epilepsy [1].

An epileptic seizure is defined as a clinical manifestation consisting of sudden and transitory abnormal phenomena which may include alterations of consciousness, motor, sensory, autonomic, or psychic events, perceived by the patient or an observer. Accordingly, epilepsy is defined as a condition characterized by recurrent (two or more) epileptic seizures, unprovoked by any immediate cause[2].

Approximately 30-40% of the patients with epilepsy have the symptomatic form, with a clear-cut brain abnormality such as a scar, a neoplasm, a vascular anomaly, or an area of cortical dysplasia. In the remaining 60-70% of cases of epilepsy, the brain appears to be normal, and the seizures are described as idiopathic [3].

Epilepsy is known from antiquity, and theories about its origin have evolved in parallel with the evolution of humankind. Thus, it is not surprising that Babylonians attributed epilepsy to magic causes[4], whereas some of our contemporaries view epilepsy as a manifestation of self-organized criticality [5].

Theoretical research in epilepsy today is boosted with the most advanced tools in neuroscience, nevertheless, the state of the art of therapies for epilepsy indicate that we are far from a full comprehension of the phenomenon: Most antiepileptic drugs and techniques appeared by chance or as result of applications of theories that were later proven wrong.

Fasting was proposed in the Bible as a way to combat demon-possessions clearly coincident with clinical pictures epileptic seizures (Mark 9:29). Ketogenic diet was introduced at the beginning of 20^th^ century as a means to mimic fasting while avoiding some negative consequences[6]. Bromides, due to their sexual desire calming effects, were proposed in 19 century to treat epilepsy [7], a malaise that according to Tissot, arises from sexual perversion[8]. Phenobarbital was given to epileptic patients as attempt to induce proper sleep [9], and valproic acid was used as a solvent for a likely anti-arrhythmic compound [10].

About 20 anticonvulsant drugs are currently available [11]. While these drugs may have side effects, such as skin rashes, nausea, sedation and memory loss, they provide seizure control in about 70% of the cases of epilepsy[12].

A brief literature review reveals that more than thousand parameters are changed in the brain as result of epilepsy presence, reflecting the extensive and complex interplay between genetic, cellular, molecular, and network mechanisms underlying the disease [13].

The diversity of observed changes accompanying epilepsy manifestations indeed illustrates the complexity of the disorder and reflects the coexistence of multiple theories about its origins and progression.

Epilepsy theories cover a broad spectrum, including hypotheses about seizure-induced brain damage, genetic mutations, neuroinflammation, network reorganization, and progressive cognitive decline. One widely discussed perspective remains Gower’s classic “seizures beget seizures” hypothesis, which postulates that seizures can progressively alter brain structure and network function, promoting further seizures and functional deterioration [14]. Yet, the evidence shows that epilepsy is not uniformly progressive—some patients achieve seizure control without decline, while others exhibit progressive cortical thinning or cognitive deficits depending on seizure type, frequency, and etiology.

Contemporary research integrates molecular, cellular, and circuit-level abnormalities alongside clinical imaging and cognitive data. There is recognition that seizure effects vary by duration, localization, and frequency, with genetic background and inter-seizure intervals modifying outcomes[15].

The plurality of epilepsy mechanistic theories today illustrates ongoing scientific efforts to frame a complex set of interacting causes and consequences, reflecting the heterogeneity of epilepsy observed in experimental models and clinical populations.

A not very epistemically sound, but goal-oriented approach in research is the Cartesian method of dividing a complicated problem into as many solvable simple components as possible. Indeed, dividing a complicated problem into many simpler, solvable components, is a goal-oriented research approach that, while not necessarily competent in capturing all complexities at once, proves highly effective in scientific inquiry, including epilepsy research. This approach supports breaking down epilepsy, a multifaceted neurological disorder, into component phenomena such as molecular mechanisms, neuronal network dynamics, and clinical manifestations, making targeted study and intervention design feasible.

In neuroscience, the Cartesian method remains foundational for studying complex systems like the brain by modeling neuronal, synaptic, or network elements separately before integrating findings to understand brain function holistically.

Among notable examples of the Cartesian method applied to neuroscience problems we mention:

- **David Marr’s Levels of Analysis** to assess brain functions at three levels: computational, algorithmic, and implementational to better understand neural computation[16].
- **Mechanistic Decomposition in Cognitive Neuroscience, in which** cognitive processes such as decision-making or sensory processing are studied by isolating contributions of specific neural circuits, neuron types, or synaptic mechanisms, analyzing each piece before integrating findings into a system-level explanation [17].
- **Modeling Brain Networks via Cartesian Grids**, allowing simpler local interpretations before considering whole-brain integration [18].
- **Decomposition of Brain Oscillations** into distinct spatio-temporal components to identify their functional significance independently, to clarify how different brain rhythms contribute to cognition and behavior[19].
- **Probabilistic Cartesian Systems** extend the classical Cartesian method to address nonlinear, stochastic brain behavior—proposing a “Nebulous Cartesian System” to account for brain-body-mind integration’s complexity, breaking down uncertain causalities systematically while acknowledging probabilistic processes [20].

In all cases, the key feature of the Cartesian method is the clear, stepwise breakdown of complex brain problems into manageable parts for analysis, followed by synthesis to form a comprehensive understanding.

This approach contrasts with, yet complements, methods focusing on emergent or network-level properties by ensuring clarity and tractability in analysis, and makes the Cartesian method a valuable, pragmatic tool in epilepsy and neuroscience research, driving hypothesis-driven experiments and computational modeling that progressively build toward comprehensive understanding despite epistemic limitations.

In this endeavor, a promising research strategy is the study of the mechanisms involved in the generation of paroxysmal depolarization shifts (PDSs), perhaps the simplest conceptual “atom of epilepsy”.

There is strong evidence that paroxysmal depolarization shifts (PDS) are a hallmark electrophysiological event in epilepsy:

- PDS represent large depolarizations in neurons that produce a burst of action potentials followed by a prolonged depolarized plateau and then hyperpolarization. This pattern is the cellular correlate of interictal epileptiform discharges seen on EEG, which are key biomarkers of epilepsy [21].
- PDS have been recorded in animal models and human epileptic tissue, with their presence associated with seizure generation[22].
- PDS underlie interictal spikes on EEG and are linked to the hyperexcitability mechanisms that generate seizures. They serve as a substrate for paroxysmal neuronal discharges characteristic of epilepsy [23].
- Electrophysiological evidence highlights their role in epileptogenesis and seizure propagation[24].

Thus, PDS are widely accepted as a fundamental cellular hallmark of epileptic activity due to their role in generating events that characterize seizures.

Being the minimal event characterizing epileptic events, hopefully, understanding PDS’s molecular and subcellular bases could help to understand the very basic elements of epilepsy.

Paroxysmal Depolarization Shifts (PDS) were first described in the 1960s and early 1970s as abnormal neuronal electrical events associated with epilepsy[21].Early pioneering studies, notably by Matsumoto and Ajmone Marsan in 1964 [25] and Prince in 1968 [26], demonstrated spontaneous PDS in cat cerebral cortex exposed to penicillin, which acted inducing epileptiform activity.

PDS are characterized by a large positive shift in neuronal membrane potential (up to +30 mV or more), lasting 40 to 400 ms or longer, during which repetitive action potentials gradually decreased in amplitude, creating a distinctive depolarizing plateau.

The discovery was significant because it identified the first cellular-level epileptiform events in an induced epileptic focus, linking membrane-level phenomena to epileptic bursts seen in EEG recordings.

Subsequent studies reproduced PDS by applying other compounds such as picrotoxin and bicuculline.

Since then, PDS has been widely studied as the fundamental electrophysiological hallmark of epileptic neurons, providing insight into seizure mechanisms and epileptogenesis, bridging clinical EEG phenomena with underlying neuronal membrane events.

A widely accepted view of PDS mechanisms is as follows:

> A paroxysmal depolarizing shift (PDS) is reflecting a giant excitatory postsynaptic potential (EPSP) due to abnormal synchronized excitatory synaptic input, often mediated by glutamate, and is linked to the generation of epileptic discharges and seizures [27-30]. PDS events can happen in clusters and are particularly associated with hyperexcitability and hypersynchrony in neuronal networks, which are essential for the emergence of epileptic spikes observed in epilepsy. Different neuronal types show variable responses to PDS, such as single or multiple action potentials during the depolarization. The PDS waveform is influenced both by the intrinsic properties of neurons and by the extracellular stimuli that trigger it. Thus, altering the neuron’s membrane potential (intrinsic property) by depolarization or hyperpolarization changes the shape and characteristics of the PDS waveform but does not change the number of PDS events in a cluster, which suggests the sequential PDS events are mainly triggered by extracellular stimuli occurring one after another rather than by a single neuron’s intrinsic activity alone [31]. Different patterns and amplitudes of stimuli produce different neuronal responses and influence PDS waveforms, which indicates that the extracellular triggering stimulus critically shapes the neuronal response and its waveform characteristics, exerting complex biophysical control over neuronal firing and PDS waveform morphology. Synaptic potentials and ionic conductances (persistent sodium currents, calcium currents) help maintain the plateau phase of the PDS, while post-PDS hyperpolarization results from calcium-activated potassium channels or GABA receptor-mediated chloride influx, showing ionic intrinsic mechanisms shape PDS waveform.
>
> Studies conclude that the PDS waveform results from complex interaction between neuron-intrinsic electrophysiological properties and the extracellular synaptic/chemical stimuli initiating the event [32].

In summary, and according to this view, the paroxysmal depolarizing shift is conceived as an abnormal neuronal electrical event signaling epileptic activity at the cellular level, involving massive excitatory input that leads to sustained depolarization and burst firing of action potentials.

The above interpretation stays in line with the widely shared view of epilepsy as a consequence of a misbalance between excitatory and inhibitory mechanisms in the brain. However, alternative views are also looming, specially supported by evidences for the possibility to obtain PDS in isolated, non-externally stimulated cells, both from invertebrates, and vertebrates’ preparations as well [33-36].

Another long-standing controversy in epilepsy research is that of epileptic neuron vs epileptic network. These hypotheses center on whether seizures originate primarily from abnormal activity in individual neurons or from pathological interactions within networks of interconnected brain regions.

**The Epileptic Neuron Hypothesis** suggests that a fixed group of damaged or dysfunctional neurons possess intrinsic properties—such as ion channel abnormalities or altered excitability—that cause them to generate uncontrolled, rhythmic epileptiform activity.

**The Epileptic Network Hypothesis.** This highly prevailing proposition posits that epilepsy is a disorder of distributed brain networks rather than single neurons or localized lesions alone. In this view, seizures arise from abnormal synchronization and dynamic interactions among multiple brain areas. The networks may involve both affected and ostensibly normal regions, suggesting that seizures reflect dysfunction in widespread circuits. This theory is supported by neuroimaging, electrophysiology, and graph theory studies that show altered connectivity patterns—even distant from lesion sites—in epilepsy patients. It also could explains why some patients without clear lesions still exhibit epilepsy and why modulation of networks (e.g., via neuromodulation therapies) can be effective.

Partisans of the epileptic neuron hypothesis recognize that the importance of synaptic events and network organization in epileptic brain is undeniable. It is obvious that even an event that originates in a single neuron, given the brain architecture, could modify the activity of a whole network, or even the whole brain. But changes observed at the network level cannot be enough to disprove the single neuron origin. Moreover, a proper discernment of the question may have implications for therapeutic strategies.

The brain’s architecture is highly interconnected and organized into complex networks with hierarchical and modular structures. This organization allows activity originating in a single neuron to rapidly influence larger neural ensembles or entire brain regions. Neural networks show rhythmic oscillations and synchronization across spatial and temporal scales, facilitating coordinated information processing, and activity from a single neuron or synaptic event is not isolated but embedded within a network context that can amplify, modulate, or gate its activity through synaptic and circuit-level interactions.

From a mechanistic perspective, this distinction is important because it means therapeutic efficacy might be achieved without addressing the root cause. This could explain why many antiepileptic drugs control seizures effectively yet do not “cure” epilepsy or reverse its molecular basis. Hence, the debate on the single neuron versus network origin has practical implications for drug development and treatment strategies.

In summary, while network-level interventions can successfully control seizure spread and improve patient symptoms, they may not reverse or modify the intrinsic neuronal defect that initiated the epileptic event. Understanding both levels—single neuron origin and network propagation—is critical for designing comprehensive therapies that target both underlying causes and delivering symptom control.

Consequently, arguments favoring the epileptic neuron view could be better supported with the study of PDS elicited under conditions where the influence from neuronal networks is minimal.

The idea of epileptic neuron has gained support from the study of epileptic activity generated in simple neuron networks, such as invertebrate ganglia. This research is devoted to the membrane pollution hypothesis (MPH), a theoretical framework proposed by Ulrich Altrup (1943-2007), centered in the epileptic neuron conception.

The membrane pollution hypothesis (MPH) [36-37] proposes that paroxysmal depolarization shifts (PDS) (the characteristic cellular hallmark of epileptic seizures [38]) arise as a consequence of aberrant pacemaker activity [39].

). According to this view, the transformation from a normal rhythmic discharge into a pathological pattern is triggered by the incorporation of amphiphilic compounds, such as convulsants or metabolic by-products, into the neuronal lipid bilayer. These intrusions alter the biophysical properties of neuronal membranes, for example by reducing membrane fluidity or, conversely, by increasing micro-viscosity.

The MPH stands in contrast to the network theory of epilepsy (NTE), which attributes seizures to synchronized hypersynchronous discharges across brain structures (particularly cortico-thalamo-cortical loops [40-41]. The NTE framework posits that epileptic activity arises from dynamic interactions among distributed regions, even when triggered by a localized seizure focus [42]. In contrast, the MPH emphasizes subcellular mechanisms, suggesting that pathological pacemaker activity embedded at the membrane level underlies seizure initiation.

Although often under-recognized in epilepsy research, several lines of indirect evidence support the MPH:

- PDS can be stably recorded from isolated nerve cells both from invertebrates [33,35], and vertebrates [34]. Certain antiepileptic drugs (AEDs) decrease membrane viscosity independently of their accepted mechanisms of action [41-49].
- Many convulsant agents reduce membrane fluidity [50-51].
- Patients with epilepsy show decreased cell membrane fluidity [52-53].
- The ketogenic diet, effective in some cases of refractory epilepsy [37], may alter membrane properties in ways consistent with MPH predictions.

Despite these findings, the MPH has not gained broad acceptance, partly because the direct causal relationship between altered membrane fluidity and epileptic discharges remains insufficiently demonstrated in humans. Epilepsy arises within the human brain, an exceptionally complex organ containing billions of neurons embedded in intricate local and global networks, where tracing straightforward cause–effect relationships is inherently difficult. Nonetheless, the fact that approximately 30% of patients remain resistant to currently available AEDs [54] highlights the urgent need for alternative conceptual frameworks that could support novel therapeutic approaches. A validated MPH could open entirely new therapeutic strategies focused on membrane stabilization, detoxification, or modulation of biophysical membrane properties, rather than the conventional targeting of ion channels or neurotransmitter systems.

To test the subcellular mechanisms proposed by the MPH, simple but informative biological models are crucial. One such system is the buccal ganglia of the snail *Helix pomatia*, which have long been recognized as well-suited for studying the cellular bases of epilepsy. Each buccal ganglion contains four large, easily identified neurons (B1–B4) on either side. These cells possess distinct physiological roles: neuron B1 acts as a “power switch” for esophageal peristalsis, B2 controls the pharyngeal salivary glands, B3 innervates the kidney, and B4 functions as a pharyngeal motor neuron [55].

Among these, neuron B3 is particularly relevant, as it reliably generates PDS that are synchronized within electrically coupled networks [35-36].

Despite their simplicity, *Helix pomatia* ganglia models closely mimic mammalian epileptiform activity. Drugs that induce seizures in humans also trigger epileptiform discharges in snail neurons [57], drugs known to aggravate epilepsy in humans enhance epileptiform activity in snails [58], and clinically effective AEDs reduce epileptiform discharges in this preparation [59]. After more than five decades of research, there is no evidence that the fundamental mechanisms of epileptogenesis in *Helix pomatia* diverge from those in vertebrates, including humans.

Crucially, in situ experiments where *Helix* neurons were perfused with Pentylenetetrazol (PTZ) revealed that PDS can be generated in single neurons, independent of synaptic activity, underscoring the role of intrinsic membrane dysfunction [60-62]. Similar results have been confirmed in cultured vertebrate neurons [63]. These findings strengthen the notion that epileptogenesis is intimately connected to the biophysical machinery underlying endogenous pacemaker potentials. What were originally described as “giant excitatory postsynaptic potentials (EPSPs)” is interpreted instead to be enlarged pacemaker potentials, justifying their description as giant pacemaker potentials. In this framework, neuron synchronization is a consequence of PDS rather than its prerequisite (likely mediated by the nonspecific release of signaling substances from epileptically active neurons into their immediate microenvironment) [64].

The observation that epileptic activity may represent aberrant pacemaker activity was first advanced by Altrup and colleagues but has not yet been directly confirmed in vertebrate models. Further theoretical support has come from Hernandez-Caceres and Brenes [56], who demonstrated through mathematical modelling that altering biophysical parameters in a Helix neuron model can transform normal pacemaker activity into epileptic discharges and eventually into sustained depolarizations.

This graded transformation unfolds through a bifurcation-like scenario, consistent with the dynamic systems perspective advanced by Lopes da Silva [65].

In light of this evidence, the present study explores whether changes in neuronal membrane micro-viscosity can mechanistically account for the transformation of a normal pacemaker potential into a PDS, thereby providing further support for the membrane pollution hypothesis.

This paper is organized as follows: The Methods section briefly describes the experiments performed to illustrate the effect of the classical convulsant PTZ on identified neurons from the *Helix pomatia* buccal ganglia, including a detailed analysis of dose dependence. Next, a version of the simple yet biophysically sound mathematical model for pacemaker activity proposed by Kononenko is presented. An effort is made to map the influence of membrane viscosity on the transition from physiological pacemaker activity to pathological paroxysmal depolarizing shifts (PDS), incorporating literature references on how increased membrane viscosity hinders conformational changes in voltage-sensitive ion channels. Specifically, increased membrane viscosity prolongs activation and inactivation time constants and shifts activation membrane potentials toward more depolarized values. A re-parametrization of the Kononenko model is proposed, representing biophysical parameters as linear functions of membrane viscosity.

The Results section begins by describing the effect of PTZ in eliciting PDS in Helix neurons. Applying the Kononenko model while increasing membrane viscosity reveals a scenario involving abrupt transitions from low-amplitude, high-frequency pacemaker activity to high-amplitude, low-frequency PDS-like activity, ultimately leading to sustained depolarizations.

The Discussion section emphasizes the theoretical possibility of simulating epileptic activity as a consequence of increased membrane viscosity. Due to the current lack of detailed experimental data supporting the details of the dependence of model’s parameters upon membrane viscosity, this study outlines the experiments needed to fill critical gaps for supporting the membrane pollution hypothesis (MPH).

## Methods

### Experimental procedure

Preparation of the buccal ganglia from the land snail Helix pomatia has been described in detail elsewhere (Altrup and Speckmann, 1994; Altrup et al., 2006). Briefly, ganglia were isolated from the animal and placed in an experimental chamber containing 1 ml of snail saline, which was continuously perfused at a rate of 3 ml/min using a peristaltic pump. Both the chamber and saline were thermostatically maintained at 20 °C. To avoid alterations in the development of epileptiform activity and to preserve the long-term viability of the preparation in situ, the ganglia were neither treated with proteolytic enzymes nor mechanically dissected to open the ganglionic capsule.

Several giant neurons were identified under a binocular microscope, allowing simultaneous intracellular recordings from multiple neurons for up to 24 hours. Glass microelectrodes filled with 150 mmol/L KCl (electrode resistance 40–50 MΩ) were employed to obtain stable measurements of membrane potential. Recordings were performed using conventional electrophysiological equipment.

The snail saline solution consisted of (in mmol/L): 130 NaCl, 4.5 KCl, 9 CaCl2, 5 Tris–Cl, with a pH range of 7.35 to 7.45. Epileptiform activity was induced by admixing the epileptogenic drug pentylenetetrazol (PTZ; Sigma, St. Louis, USA) to the saline. Concentrations of PTZ applied included subthreshold doses (1, 4, 8, and 16 mmol/L) as well as reliably effective suprathreshold concentrations of 40 and 80 mmol/L. Membrane potential changes were recorded via an ink recorder set with a time constant of approximately 5 ms, sufficient to capture both pacemaker potentials and paroxysmal depolarizing shifts (PDS).

All animal procedures complied with relevant international, national, and institutional guidelines. Efforts were made to minimize animal use and suffering. Experiments adhered to Directive 2010/63/EU of the European Parliament and Council on the protection of animals used for scientific purposes (22 September 2010).

### Representation of minimal Kononenko’s model (MKM)

In 1994, Kononenko [66] proposed a model, inspired by the Hodgkin-Huxley framework, to represent pacemaker activity in the bursting neuron RPa1 of *Helix pomatia*. This is one of the simplest biophysically sound pacemaker models available, and successfully preserves the core features of a pacemaker generator. We refer to it as the “Minimal Kononenko Model” (MKM).

#### The MKM includes the following components

- A linear leak pathway characterized by voltage-independent sodium (g_Na) and potassium (g_K) conductances.
- A persistent, voltage-dependent, stationary sodium conductance (gNa_V), accounting for the negative resistance region observed in the stationary current-voltage relationship of bursting neurons.
- A time-dependent conductance (g_B) activated by hyperpolarization, regulated by Hodgkin-Huxley-type kinetics involving activation and inactivation variables ‘m’ and ‘h’.

The MKM effectively replicates a range of experimentally observed phenomena in the RPa1 neuron, such as an increase in input resistance during the interburst interval, the oscillation period’s dependence on polarizing current, modulation of burst activity, and behavior of ionic currents under boundary voltage conditions. Originating from rigorous electrophysiological research, the MKM serves as a robust foundation for subsequent studies.

Moreover, the MKM and its variants have revealed diverse dynamic regimes, including chaotic bursting patterns, providing insights into the complex behavior of pacemaker neurons [67-69].

#### The MKM takes the form

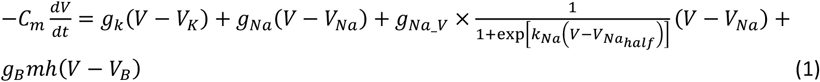

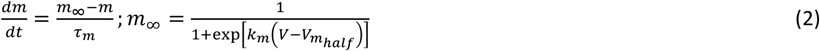

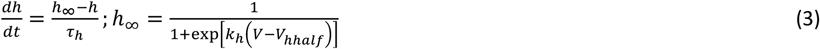

In equations (1–3), Cm represents the membrane capacitance in µF, while V denotes the membrane potential in mV. The variables m and h are the voltage-dependent Hodgkin-Huxley activation and inactivation gating variables for the gB conductance respectively. The parameters k_m_ and k_h_ characterize the voltage dependence of these gating variables; V_*mhalf*_ and V_*hhalf*_ correspond to the voltage at which half of the corresponding particles are at the favorable position.

Even when simple, MKM incorporates a total of 18 parameters, each with a well-defined and consistent biophysical meaning [66].

### Modelling the effect of viscosity on pacemaker potential using the MKM

Membrane viscosity is a significant biophysical factor influencing the gating kinetics of voltage-gated ion channels, particularly the activation (m) and inactivation (h) particles in excitable membranes. This adds a critical layer to the Hodgkin-Huxley framework, where gating is not solely voltage-dependent but also sensitive to the membrane’s physical state.

Evidence suggests viscosity modulates ion channel function by affecting lateral mobility and conformational dynamics. For instance:

* Heine et al. (2016) review how viscosity and thermal energy modulate channel mobility and flexibility [70].
* Kashirina et al. (2020) used molecular rotors to demonstrate that fluidity changes affect channel behavior in stem cells [71].
* Almog, Degani-Katzav, and Korngreen (2022) provide kinetic modeling evidence that viscosity alters m and h particle dynamics [72].
* Different evidences showed that lipid composition changes, which alter viscosity, affect voltage gating [73-75].

The primary mechanism involves higher viscosity physically hindering the movement of voltage-sensing domains (S4 helices), thereby delaying channel opening and potentially altering the energy required for conformational changes. Supporting this, molecular dynamics simulations show reduced S4 segment mobility in viscous membranes [76], whereas PIP_2_ signaling lipids can reduce local viscosity, accelerating activation [77].

The effects on inactivation are also documented. For example, n-3 PUFAs can shift inactivation voltage dependence and alter recovery rates [78-79], with similar effects observed on calcium currents [80-81]. These effects may share a mechanism with general anesthetics, which are thought to act via alterations in membrane fluidity [82].

Given that MKM model parameters are quantitatively dependent on membrane viscosity, and lacking specific data for snail membranes, we focused on the well-documented effects on classical voltage-dependent channels particle kinetics modification by membrane viscosity. Key quantitative findings include:

* Kinetics: Benzyl alcohol (decreasing viscosity) increases the activation rate constant (α_m_) by 15-25% [83], correlating to a ∼5% viscosity change [84]. A 10°C temperature increase (reducing viscosity) decreases the activation time constant (τ_m_) by a factor of 2-3 [85].
* Voltage Dependence: Cholesterol (increasing viscosity) causes a +10 to +15 mV shift in the half-activation voltage (V_1_/_2_) [86], corresponding to a fourfold increase in microviscosity index [87]. Conversely, long-chain PUFAs (decreasing viscosity) shift V_1_/_2_ by approximately −7.5 mV [88].

In summary, membrane fluidity acts as a powerful allosteric regulator, directly modulating the energetics and kinetics of the voltage-sensing particle’s movement, providing a basis for its integration into the MKM model.

In order to take into account, the modulation by membrane viscosity of relevant biophysical events in voltage-dependent excitability, we reparametrized equations 1-3, taking into account the following relations respect to membrane viscosity:

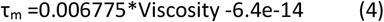

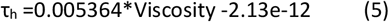

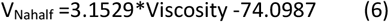

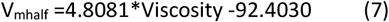

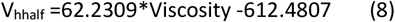

The initial values were selected as represented in Table 1. All values are close to those reported by previous studies [65-66, 86-88]. The value of viscosity (8.85 cP) is in the range of reported figures for living cells (8 to 24 centiPoise. cP), [89].

**Table 1.** Initial values for the modified MKM (equations 1-6). In the implemented Matlab simulation, the viscosity parameter was changed in steps of 0.01 cP.

|  |  |
| --- | --- |
| Viscosity0 | 8.85 cP |
| V(1) | -50 mV |
| hb(1) | 0.77 |
| mb(1) | 0.54 |
| Cm(1) | 0.03 |
| I | 0.0 nA |
| Ena | 40.0 mV |
| EK | -61.0 mV |
| Eb | -76 mV |
| gNa | 0.0556 |
| gK | 0.22 |
| gB | 0.5 |
| gNa_V | 0.146 |
| Vnhalf | -24.99 mV |
| kNa | -0.30 |
| khb | -0.55 |
| kmb | 0.4 |
| VNahalf | -45.9 mV |
| Vhbhalf | -55.9 mV |
| Vmbhalf | -49.4 mV |
| Taumb | 0.025 s |
| Tauhb | 1.9 s |

## Results

### Recording epileptic activity in snail buccal ganglia

The response to PTZ application (40 mM) by neurones B3 and B4 is illustrated in figure 1.

**Figure 1.**
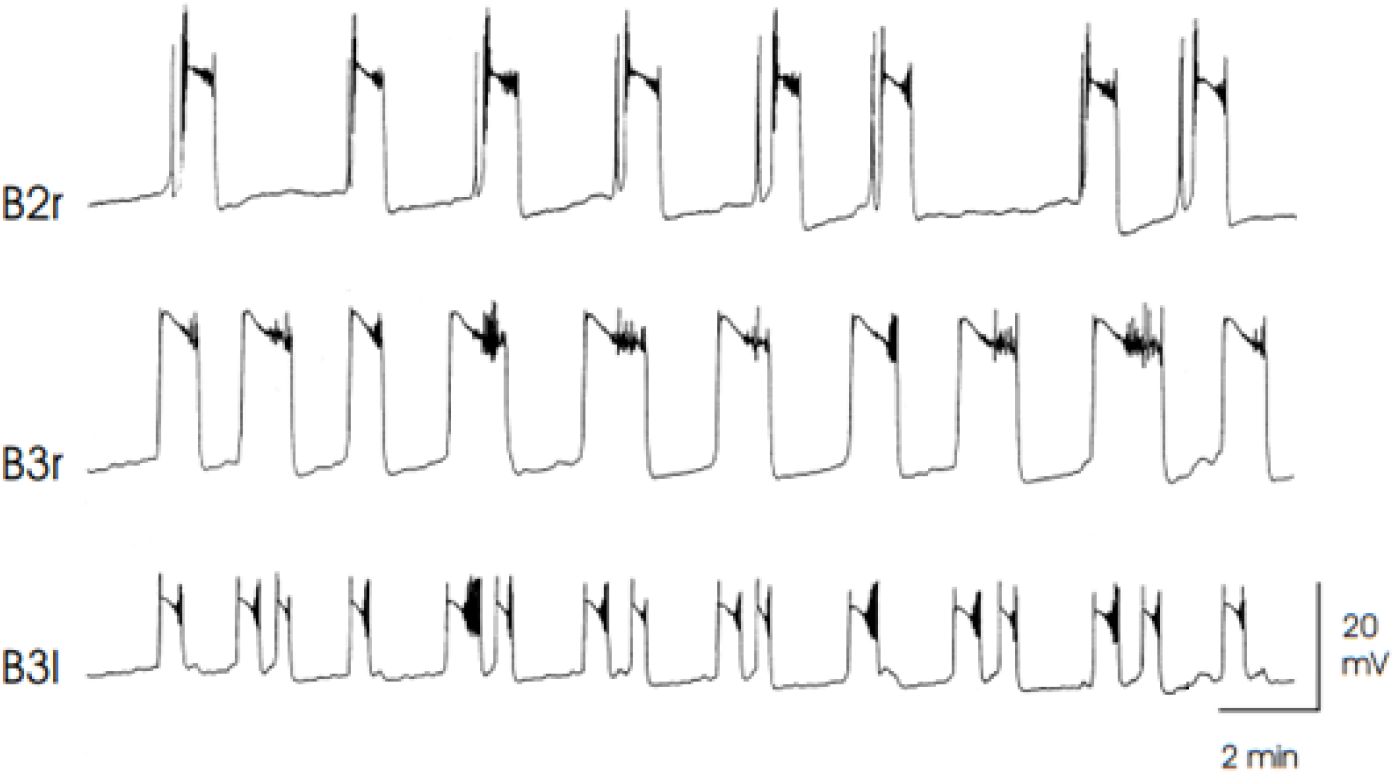
Paroxysmal depolarization shifts observed in three simultaneously recorded nerve cells. Only the right buccal ganglion was perfused with PTZ (40 mM).

The similarity and a certain degree of synchrony are apparent, in spite of their differences in physiological function [35], as well as the fact that only one hemiganglion was perfused.

In this preparation, the PDS developed at the right ganglion’s B3 neuron (B3r) are setting the pace. In B3r PDSs have the highest amplitude and longest duration, and they precede in the time axis the appearance of epileptic activity in B2r. Precedence in B3 is not very evident, perhaps due to the electrical coupling between both B3 neurons [35] 35.

PTZ exerts its effect in a dose dependent way. This is illustrated in figure 2. Unlike typical dose-effect relationships. We observe that pentylenetetrazol induces quantitative changes on the electric activity of the cell. Whereas 4 mM induces the appearance of pacemaker activity, which steadily increases up to 16 mM, at 40 mM typical, full developed PDS appear. At 80 mM PDS evolve into sustained depolarizations, perhaps reflecting the cellular electrophysiological bases for status epilepticus or generalized seizures.

**Fig. 2.**
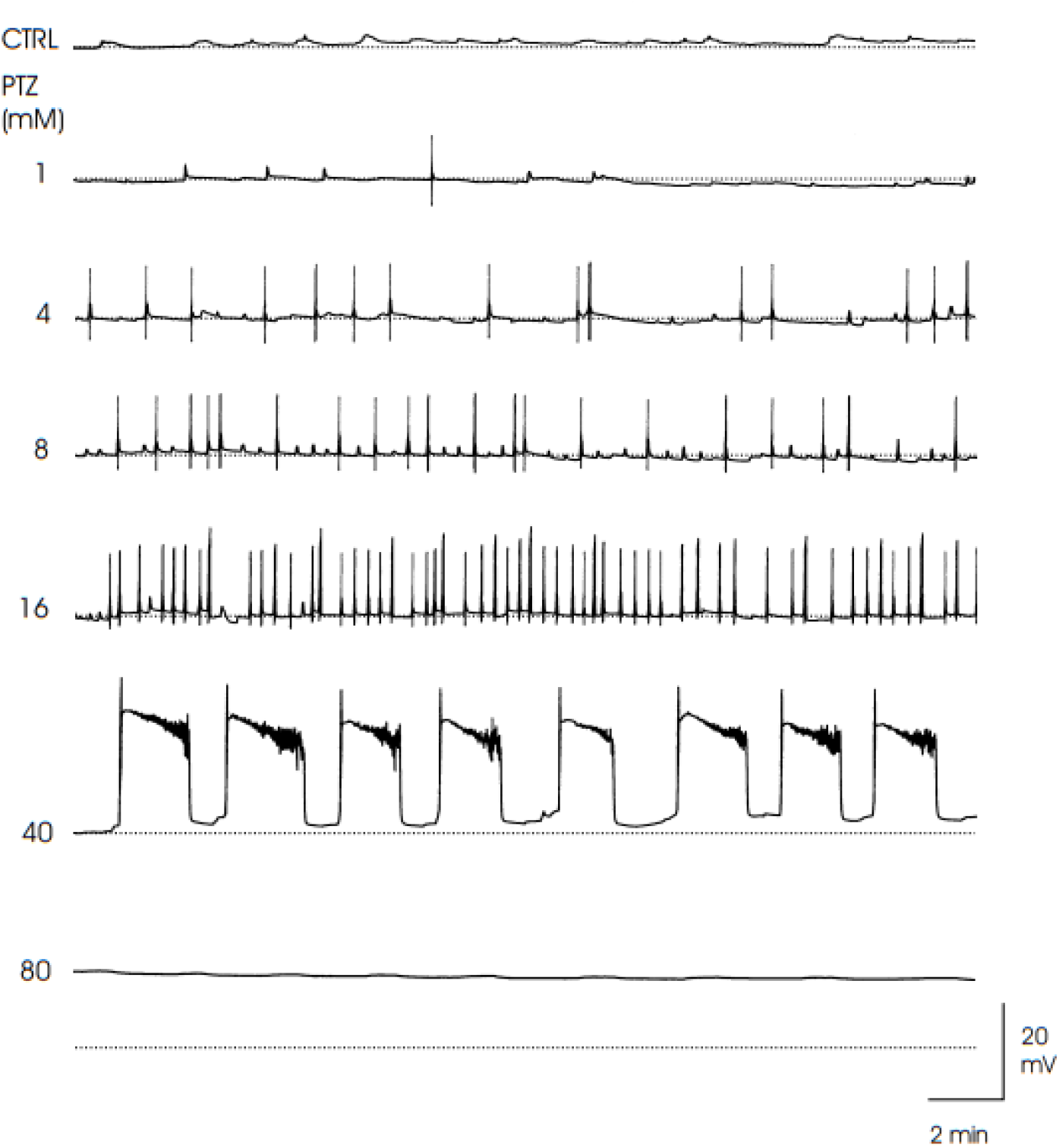
Effect of increasing PTZ doses on neuron B3. Ink recording of buccal ganglia neuron B3 activity without stimulation (control condition, CTRL) and under treatment with different concentrations (8-80 mM) of the epileptogenic drug pentylenetetrazol (PTZ). The dotted line represents the original baseline, close to the resting membrane potential at −50 mV. In particular, it was shown that this scenario is unrelated to synaptic mechanisms, [35]. As Altrup et al (2004) state, “PDS may be labelled as “giant pacemaker potentials” which are generated by the single neurons”[38].

This pattern is typical for other convulsants acting on neuron by, such as etomidate [35] and penicillin [90].

Altrup and others [35] remarked that the fact that chemically very different substances evoke an identical pattern of responses on neurone B3 should suggest of a common mechanism.

#### Simulating the effect of membrane viscosity upon the transformation of pacemaker potentials upon epileptic activity

Our further step was to explore the possibility to obtain PDS-like activity in a model for pacemaker potentials from Helix neurons. In a recent study, Hernandez-Cáceres and Brenes [56] showed that modifying the parameters of the MKM (equations 1-3) it is possible to mimic such transformation.

Here we are using a modification that takes into account the influence of membrane viscosity upon MKM parameters (equations 4-6), and using the fixed initial values set in Table 1. As shown in figure 3 (A-E), solutions mimicking stable membrane potential (3A), low-amplitude pacemaker activity (3-B), Higher amplitude pacemaker activity (3-C), low frequency potentials resembling PDS (3-D) and sustained, long-lasting depolarizations.,

**Figure 2A.**
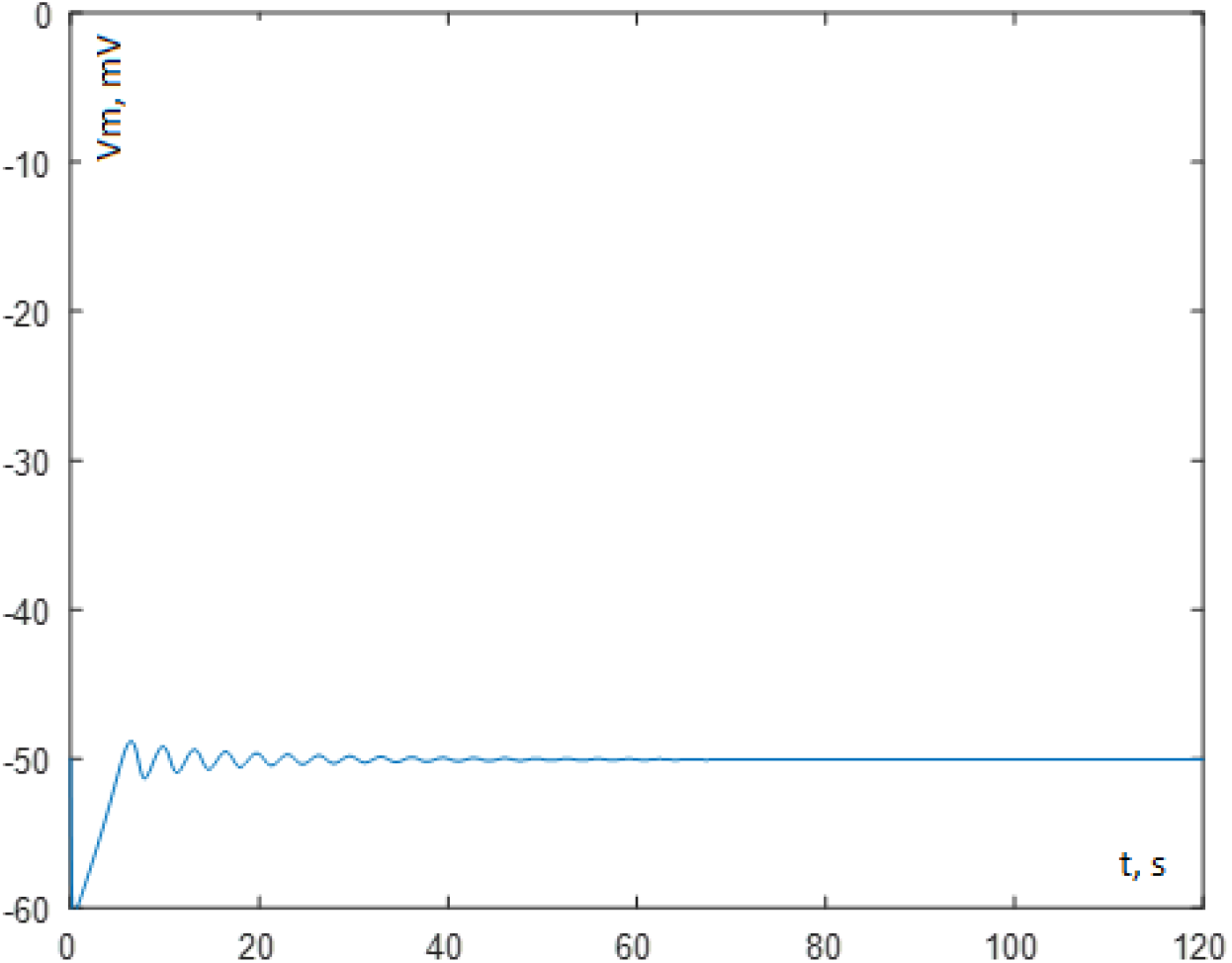
Damped oscillations corresponding to the modified MKM with micro-viscosity= 9.04 cP.

**Figure 2B.**
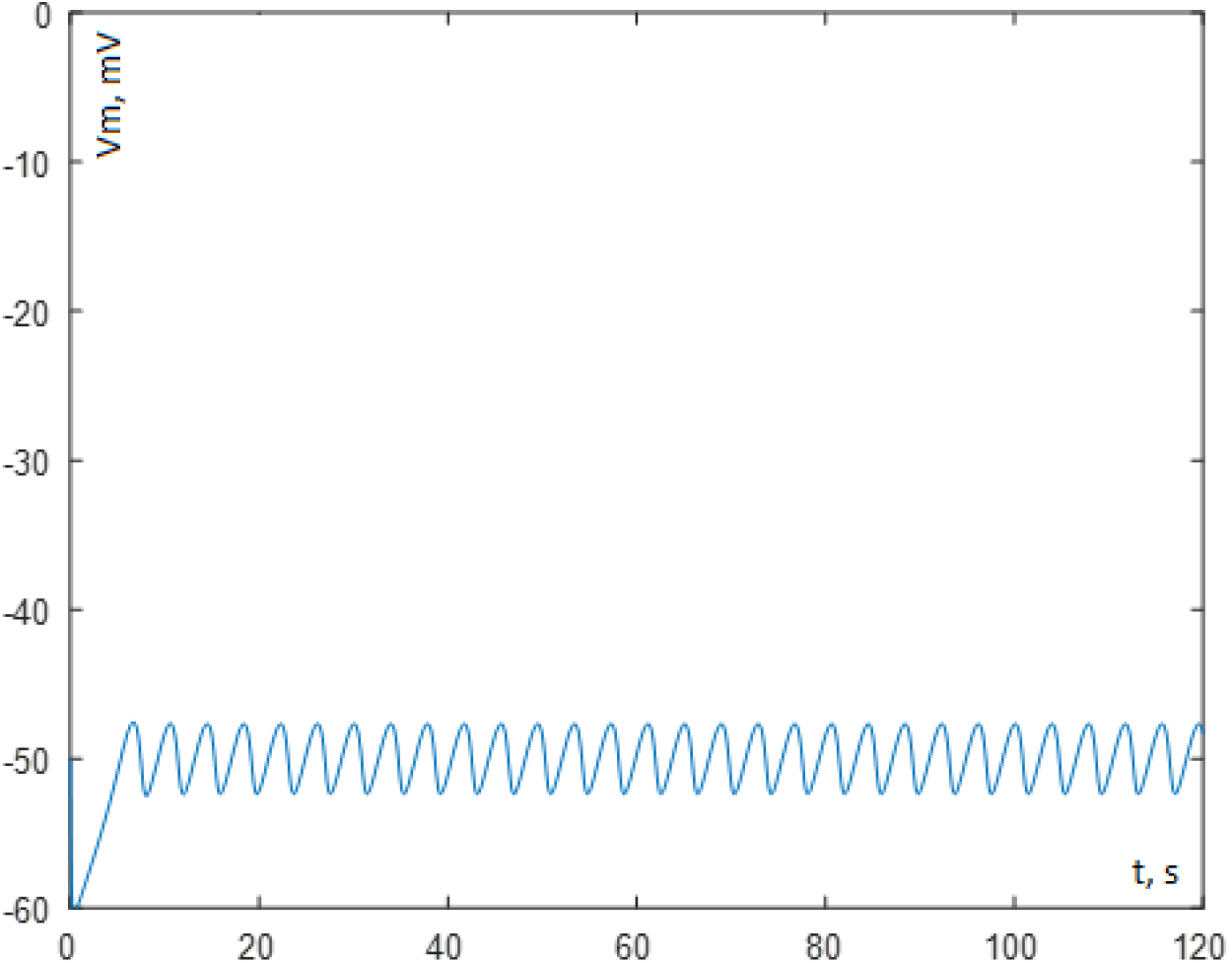
Pacemaker activity obtained with the modified MKM with micro-viscosity= 9.28 cP.

**Figure 2C.**
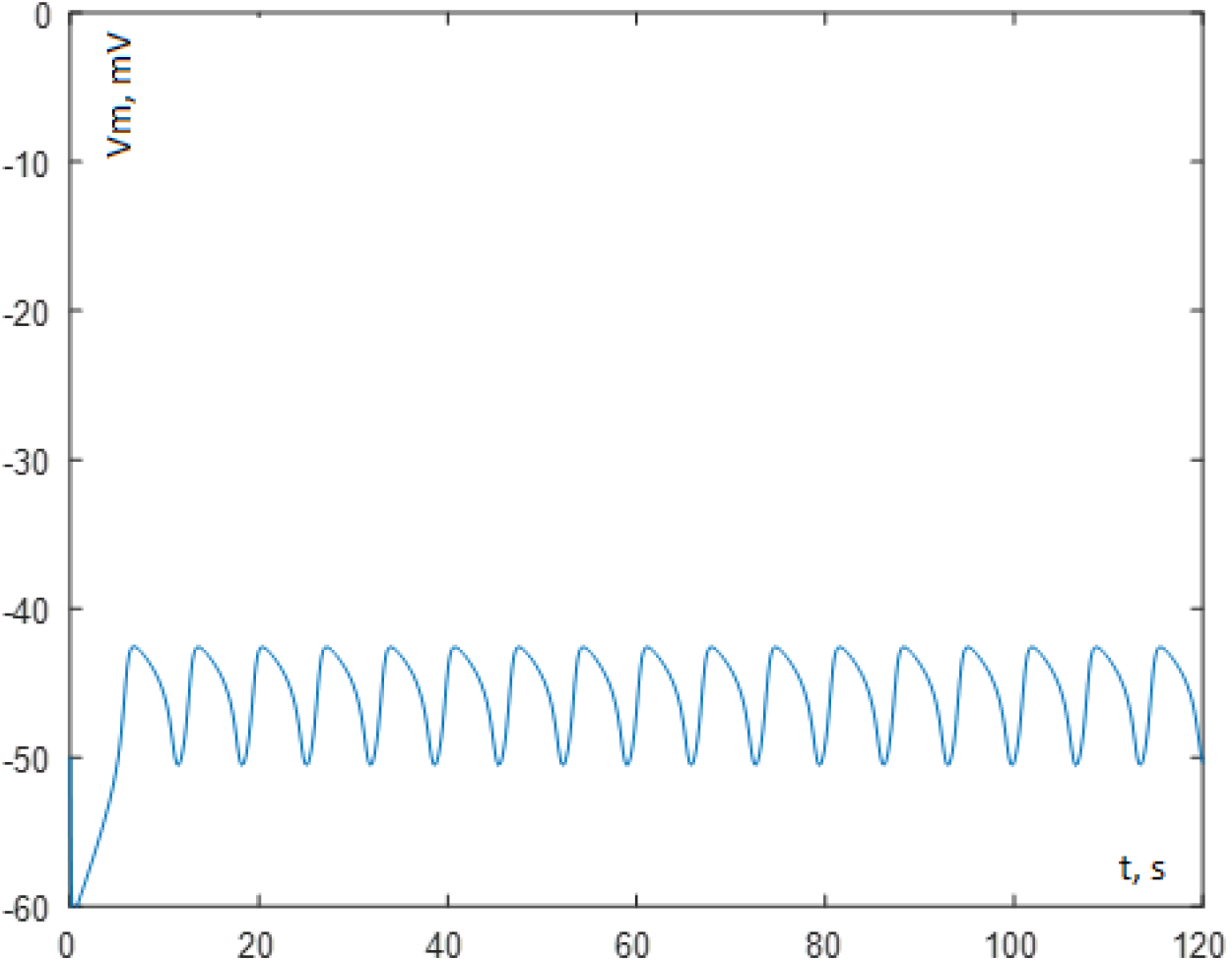
PDS generated with the modified MKM with micro-viscosity= 9.63 cP.

**Figure 2D.**
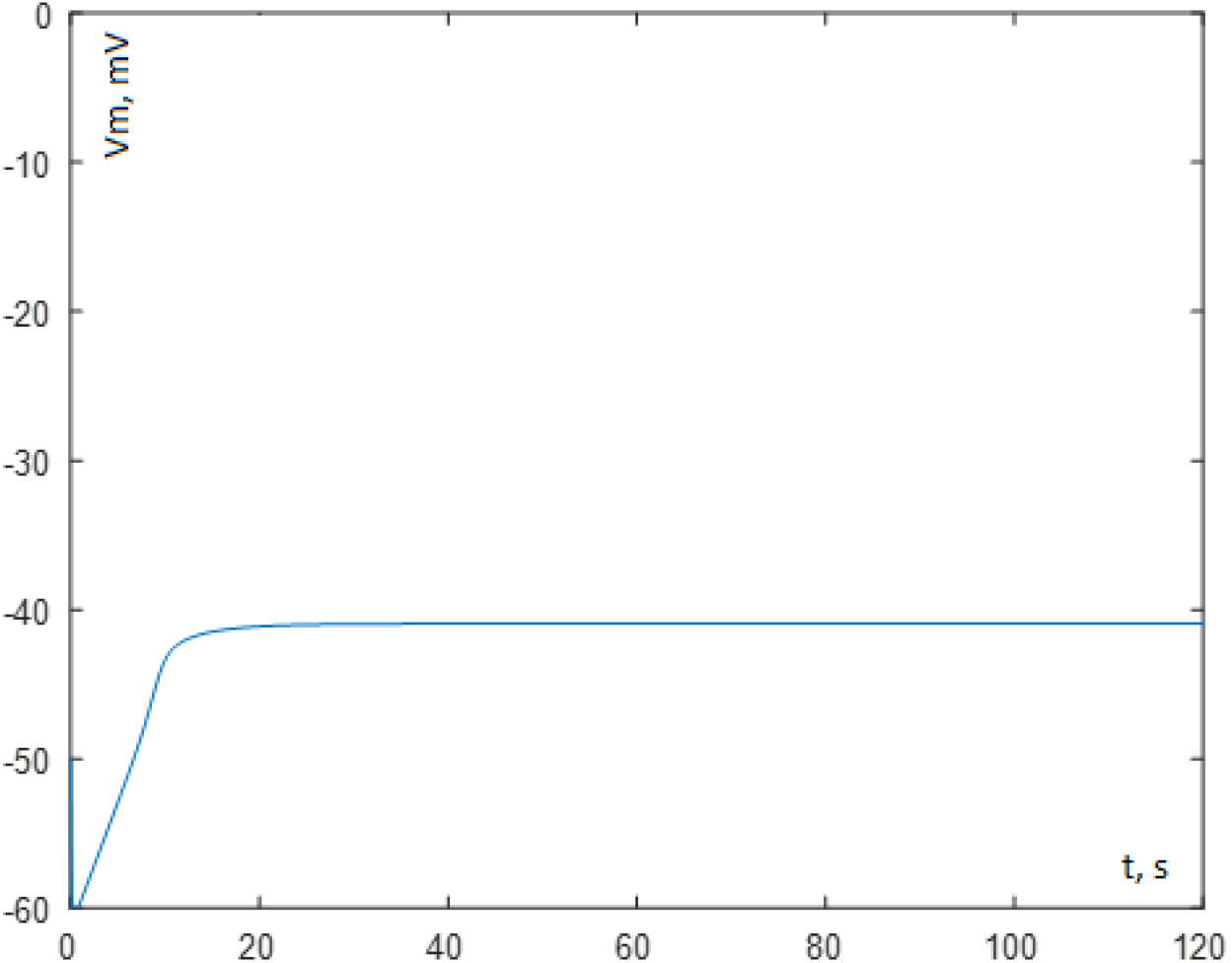
Long-lasting depolarization corresponding to micro-viscosity= 9.854 cP. This could correspond to generalized seizures or status epilepticus.

By varying the viscosity parameter in small steps, we obtained the results represented in figure 4. As apparent, the behavior from one type of activity into another might proceed through a bifurcation scenario: little variations in the viscosity parameter move the solution from stable resting membrane potential to oscillations of increasing amplitudes, and further to PDS-like behavior, and finally to sustained depolarizations.

**Figure 4.**
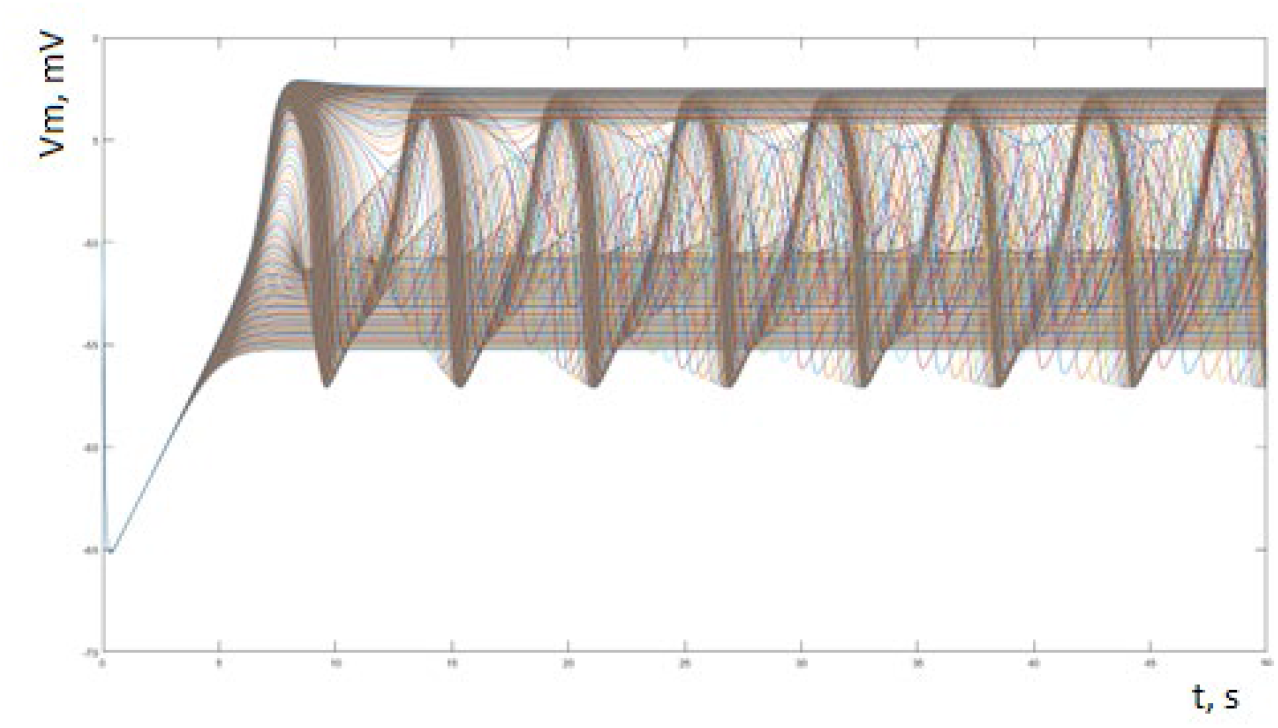
Superposition of modified MKM solutions by varying the viscosity parameter from 8.85 cP to 9.85 cP in steps of 0.01 cP.

## Relating experimentally recorded behavior to membrane viscosity

At this stage, no direct experiments were performed measuring membrane viscosity of snail neurons under different PTZ concentrations. The only indirect information is available from the measurement of stationary lateral pressure of lipid monolayers at different PTZ concentrations (and Etomidate) (figures 9 A-B from [1]).

The quantitative relationship between monolayer lateral pressure (surface pressure, π) and membrane (monolayer) surface viscosity (η) has been studied, and a linear dependence of viscosity respect lateral pressure is a good empirical approximation for the range of lateral pressures measured by Altrup et al [1].

Combining empirical results from ref [91] and ref [92].

We are suggesting the following linear relationship.

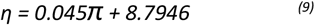

this provides the relationship between both variables that is represented in figure 5.

**Figure 5.**
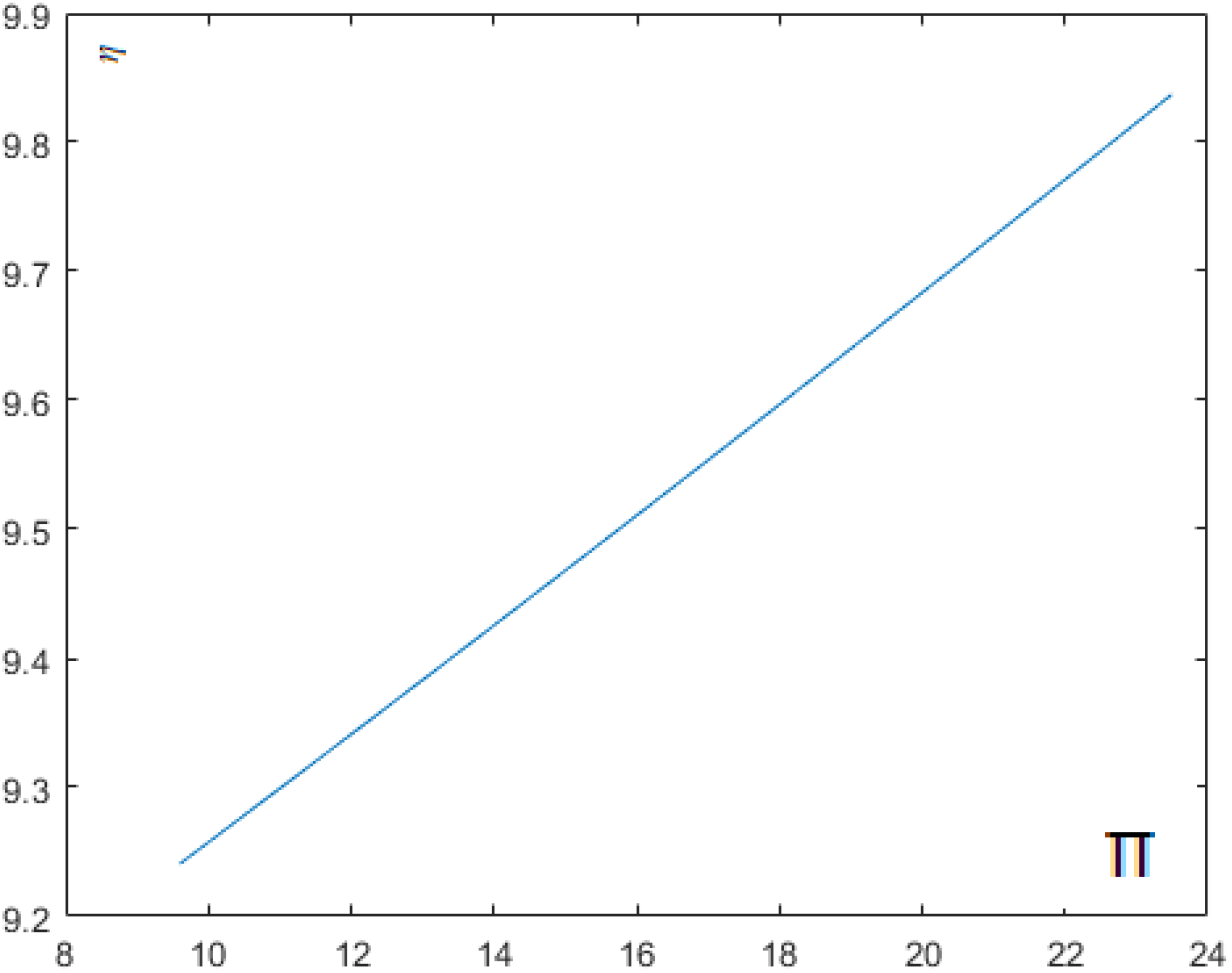
relation between lateral pressure π (x-axis) and the viscosity parameter *η* (y-axis) according to Equation (9).

**Table 2.** Suggested micro-viscosity values corresponding to different PTZ concentrations. Values for lateral pressure π were obtained from experimental data from reference [1] (Figure 9). Microviscosity *η* was calculated from equation (10).

| [PTZ]<br>mM | $\pi$ mN/m | $\eta$ cP |
| --- | --- | --- |
| 1 | 9.7 | 9.22 |
| 4 | 11.2 | 9.30 |
| 8 | 11.4 | 9.32 |
| 16 | 12.8 | 9.38 |
| 40 | 16.7 | 9.55 |
| 80 | 23.6 | 9.85 |

With this transformation, we can obtain a comparison between experimental and simulated results, as it is represented in figure 6

**Figure 6.**
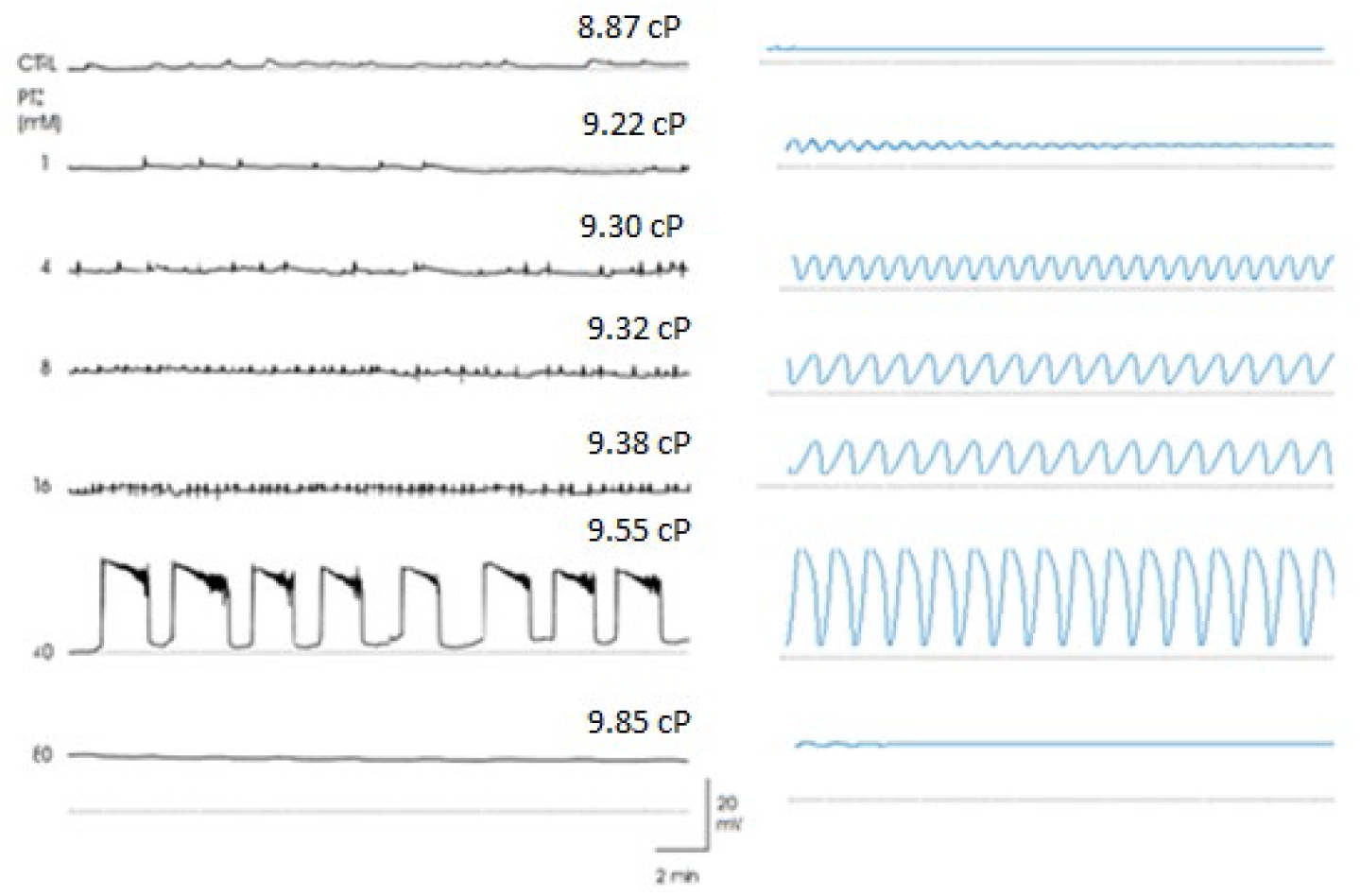
comparison between experimental (left panel) and theoretical (right panel) results for the effect of PTZ on neuron B3 (figure2)

As noticed, good agreement between experimental and theoretical results has been obtained.

## Discussion

This study aims to demonstrate the plausibility of the membrane pollution hypothesis (MPH), a theoretical framework that faded from scientific discourse following the unexpected death of Ulrich Altrup in 2007. The hypothesis has suffered from limited experimental evidence, a lack of further theoretical development, and its divergence from mainstream views in epileptology. However, the existence of a physiologically sound model for pacemaker activity, prior work showing that epileptic activity may arise from pacemaker potentials[22] 56, and experimental and theoretical evidence that membrane viscosity can alter key biophysical parameters, in particular those included in the MKM collectively suggest a credible scenario. This scenario illustrates how changes in membrane viscosity may “lead to the expression of the membrane pacemaker machinery”[1] and induce paroxysmal depolarizing shifts (PDS) as well as sustained depolarizations. A notable finding is that the transition from pacemaker activity with increasing membrane fluidity follows a bifurcation-like pattern, consistent with the concept of epilepsy as a dynamical disease [32].

Experience in biosciences shows that enduring models are those grounded in rigorous experimental protocols and quantitative measurement of relevant biophysical parameters. This is exemplified by Mendel’s mathematical inheritance model for *Pisum*, based on extensive experimental work over many years, and by the Hodgkin-Huxley model, which was strongly supported by detailed voltage clamp experiments on squid giant axons. Similarly, the MKM is founded on more than a decade of careful experimental research by N.I. Kononenko. Unfortunately, our current study lacks direct experimental assessment of how changes in membrane microviscosity affect the specific parameters measured by Kononenko. Therefore, the proposed quantitative relationships (Equations 4–8) relating these parameters to micro-viscosity, as well as the expression (Equation 10) linking membrane viscosity to lateral pressure, remain theoretical and unconfirmed experimentally. Nonetheless, our results provide a plausible theoretical framework supporting the core tenets of the MPH.

## Limitations

In this study, viscosity values were derived from plausible assumptions rather than direct measurements. Additionally, the impact of viscosity changes on the time constants and half-activation potentials for the ion currents described in the MKM has yet to be empirically characterized. Consequently, our findings propose a qualitative scenario rather than providing precise quantitative predictions. We anticipate that with further experimental data, the modified MKM can be refined, enhancing its predictive power and enabling extrapolation of our key conclusions to epileptogenesis in the mammalian brain.

## Conclusion

Our results lend additional theoretical support to the MPH proposition that increases in membrane microviscosity can transform normal physiological pacemaker potentials into pathological paroxysmal depolarizations.

## Acknowledgements

This research was supported by the Cuban Program for Neuroscience and Neurotechnology (PN305LH013-070), as well as the project № Б23КУБ-026 supported by the Belarussian Fond Supporting Fundamental Research (БРФФИ). Authors express their gratitude to Prof. Oscar Brenes, from the University of Costa Rica for encouragement and useful suggestions.

